# Accidental and Regulated Cell Death in Yeast Colony Biofilms

**DOI:** 10.1101/2025.02.02.636168

**Authors:** Daniel J. Netherwood, Alexander K. Y. Tam, Campbell W. Gourlay, Tea Knežević, Jennifer M. Gardner, Vladimir Jiranek, Benjamin J. Binder, J. Edward F. Green

## Abstract

The yeast species *Saccharomyces cerevisiae* is one of the most intensively studied organisms on the planet due to being an excellent eukaryotic model organism in molecular and cell biology. In this work, we investigate the growth and morphology of yeast colony biofilms, where proliferating yeast cells reside within a self-produced extracellular matrix. This research area has garnered significant scientific interest due to its applicability in the biological and biomedical sectors. A central feature of yeast colony biofilm expansion is cellular demise, which is onset by one of two independent mechanisms: either accidental cell death (ACD) or regulated cell death (RCD). In this article, we generalise a recently developed continuum model for the nutrient-limited growth of a yeast colony biofilm to include the effects of ACD and RCD. This new model involves a system of four coupled nonlinear reaction–diffusion equations for the yeast-cell density, the nutrient concentration, and two species of dead cells. Numerical solutions of the spatially one and two-dimensional governing equations reveal the impact that ACD and RCD have on expansion speed, morphology and cell distribution within the colony biofilm. Our results show good qualitative agreement with our own experiments.

## 1 Introduction

We investigate how two nutrient-dependent cell death mechanisms affect the expansion speed, morphology, and cell distribution of yeast colony biofilms. Biofilms are communities of microorganisms that reside within a self-produced extracellular matrix (ECM) [1], which itself is adhered to a surface. The ECM protects the biofilm from external insults [2, 3] and facilitates efficient transportation of nutrients and water [4]. More than 80% of all microbial life is found within biofilms [5], making them one of the most prevalent life forms on Earth [3]. Biofilms impact human life in several ways. Positive impacts include: wastewater treatment, food preservation, and other areas of biotechnology. However, the main impact of biofilms is their role in pathogenic bacterial and fungal infections [5]. Biofilms are known to colonise medical devices, and are a leading cause of hospital-acquired infections. Yeast biofilms of *Candida albicans* yeasts [6] cause invasive candidiasis, a disease responsible for approximately 20% of bloodstream infections in intensive-care units [7]. These major impacts, together with the rise of antimicrobial resistance, motivate the continued research into the mechanisms that promote and inhibit biofilm expansion.

In this work, we consider colony biofilms formed by the yeast species *Saccharomyces cerevisiae*, which is canonically referred to as the baker’s yeast. *S. cerevisiae* is a major model organism in biofilm research [8–11], and was the first eukaryote to have its genome sequenced [12]. Soon after, Reynolds and Fink [8] demonstrated that *S. cerevisiae* can form colony biofilms on semi-solid agar. These colonies are single-species structures consisting of cells and ECM. They represent an experimental model for the formation of more complicated multi-species yeast biofilms found in medical and industrial settings [13, 14]. Throughout this work, we use the terms colony biofilms or colonies to describe these single-species entities, in accordance with Plocek et al. [15].

Yeast species can alter their growth in response to their environment. For example, on the microscopic scale, individual yeast cells are thought to forage or move away from the colony by elongating, a process that creates filamentous patterns [16]. On the macroscale, entire colonies are known to develop nutrient-driven spatiotemporal instabilities, leading to floral morphologies [14, 17]. In this work, we investigate how cell death affects the growth and dynamics of colony biofilms. Figure 1.1 illustrates three examples of *S. cerevisiae* colony biofilm growth from our own experiments. In each experiment, the yeast colony is stained with a water-soluble red dye, Phloxine B [18–21]. An increased intensity of red colour in each photograph is indicative of regions within the colony where membrane permeability has been compromised due to a sufficient drop in metabolism, which is a strong indication of cell death [22]. In Figure 1.1A, there is a large circular region of cell death in the centre of the colony, surrounded by mostly living cells close to the leading edge. This pattern resembles that of a necrotic core observed in spheroidal tumour growth [23–25]. In Figure 1.1B, the colony expands axisymmetrically, having a distinct red ring of cell death trailing the leading edge. There is also a region of elevated cell death near the centre of the colony, albeit not as pronounced as the ring or in the core of Figure 1.1A. Figure 1.1C shows a colony biofilm grown in a rectangular Petri dish, which also shows a stripe of dead cells trailing the edge of the colony. The colony in Figure 1.1C was initiated as a thin stripe of cells in the centre. During the expansion, the colony develops a spatially non-uniform front shape, similar to the petal formation in circular experiments reported by [14]. Here, we investigate how nutrient limitation and cell death influence the different cell-death patterns and colony morphologies observed in Figure 1.1.

**Fig. 1.1:**
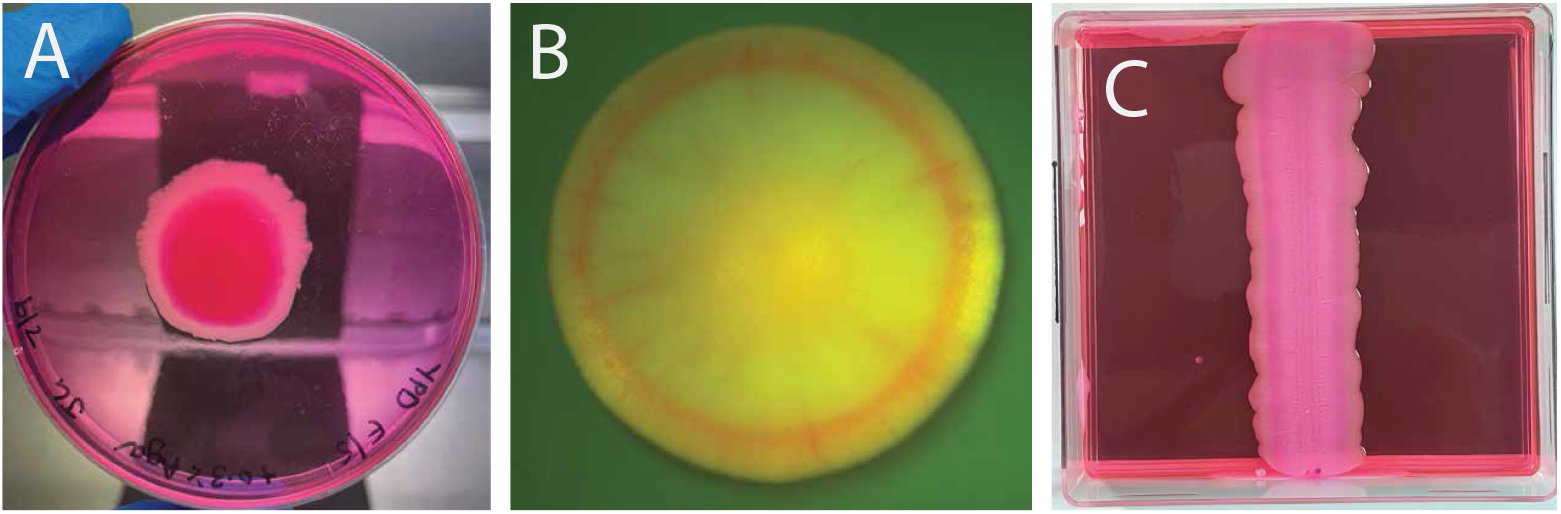
Three experimental photographs of yeast colony biofilm formation. Dark pink/red regions in each photograph shows regions where yeast cells have taken up Phloxine B dye, indicating regions of cell death. The experimental method is described in Section 2.2. (A) *Magnusiomyces magnusii* colony biofilm grown from a small spot inoculation at the centre of a circular Petri dish (10 days), with cell death occurring in the centre of the colony. (B) *S. cerevisiae* colony biofilm grown on agar (5 days). (C) *S. cerevisiae* colony biofilm grown from an inoculum streaked from top to bottom of a rectangular Petri dish filled with 0.9% YPD agar (19 days), with a stripe of death trailing the expanding edge, and an unstable front pattern.

The review by Carmona-Gutierrez et al. [26] summarises the current research in yeast-cell death, highlighting two specific mechanisms of cellular demise, namely Accidental Cell Death (ACD) and Regulated Cell Death (RCD). ACD occurs when cells encounter harsh and unpredictable environments, such as insufficient nutrient supplies [27]. This cell-death mechanism results in a necrotic cell morphotype, whereby cells experience structural disintegration and uncontrolled rupturing of the plasma membrane. On the other hand, RCD is cellular suicide designed to benefit the survival of the colony as a whole. RCD typically occurs in response to mild stresses, though it can be entirely initiated by the cell. RCD yields a spectrum of cell-death morphotypes. An important hypothesis is that RCD can release nutrients back to the surrounding environment for further consumption by the cells [28, 29]. RCD can therefore have a positive impact on the colony by producing an additional nutrient supply, promoting growth.

Whilst there have been recent advancements in measuring the lifespan of a cell [30, 31], obtaining an explicit diagnosis of cell death remains an experimental challenge. This is because a diagnosis of cell death can be made only when irreversible breakdown of the plasma membrane or complete cellular fragmentation is detected [26]. The vitality dye Phloxine B used in our experiments is only retained in metabolically inactive cells that cannot expel the dye. Whilst this is a reliable indication of cell death, it does not explicitly measure death, nor the mechanism by which the cell may have died. For this reason, we turn to mathematical modelling for further insight. Scientists have used agent-based [32–34], hybrid [35], reaction–diffusion [14, 17, 36–40], and mechanical [41– 48] mathematical models to study biofilm formation and growth [49, 50]. Work has been done to incorporate cell death into some of these models [35, 51, 52]. Ghosh and Levine [35] used an agent-based model and reaction–diffusion system with a Heaviside step function in the reaction term to show that nutrient-limited cell death (an instance of ACD) and cell death occurring randomly throughout the colony both amplify front patterns. Whilst clear progress has being made to improve our understanding of the impact that cell death has on colony biofilm expansion, comparatively little is understood about the effect that the mechanisms of ACD and RCD, and their breakdown and subsequent nutrient release have on the growth dynamics.

Here, we develop a continuum reaction–diffusion model incorporating nutrient limitation, ACD, and RCD, as well as cellular breakdown and nutrient release. We neglect mechanics and the ECM, and start from a system of nonlinear reaction–diffusion equations for nutrient-limited growth considered by several authors [14, 17, 36], and modify the model to incorporate cell death. This original model investigates a colony biofilm occupying a fixed Petri dish, and involves the following system of reaction–diffusion equations:

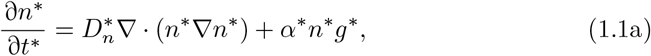

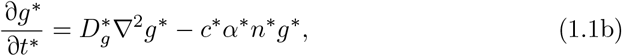

where *n*^*^(***x***, *t*) and *g*^*^(***x***, *t*) are the dimensional living-cell density and nutrient concentration at dimensional time *t*^*^ and dimensional position ***x***^*^. Since yeast-cell diffusion occurs more rapidly in regions of elevated cell density, living-cell diffusion is inherently nonlinear [36]. We therefore adopt the nonlinear degenerative diffusion law used previously by Müller and van Saarloos [36] and later by Tam et al. [14], where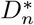 is the dimensional living-cell diffusivity constant. In this model, nutrients are assumed to diffuse according to Fick’s law with dimensional diffusivity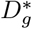. The colony expands through cell proliferation, facilitated by nutrient consumption. The constant *α*^*^ is the dimensional rate of cell proliferation, and *c*^*^ is a dimensional constant indicating the amount of nutrient required to produce a new cell.

A common technique applied to reaction–diffusion systems such as (1.1) is to analyse the spatiotemporal stability of transversely-perturbed travelling-wave solutions [53– 57]. As demonstrated by Tam et al. [14] and others, the minimal reaction–diffusion model (1.1) can be used to predict the expansion speed and front patterns of yeast colony biofilms in the absence of cell death. In this article, we introduce ACD and RCD into (1.1) and investigate numerically their influence on the systems evolution into a non-uniform front shape.

The remainder of the paper is organised as follows. In Section 2, we present the mathematical model and experimental method. We use a four-species reaction–diffusion system for yeast-cell density, nutrient distribution, and the density of cells having undergone either ACD or RCD. We assume that ACD occurs when the nutrient concentration drops below a prescribed threshold, and that RCD occurs in a prescribed window of nutrient concentration. In Section 3.1, we show that our model is capable of producing the patterning observed in Figure 1.1. In Section 3.2, we demonstrate qualitatively that cell death in isolation amplifies the instability of a planar front, causing a more pronounced petal formation. Our results indicate that RCD can offer a survival advantage to the colony, increasing expansion speed, and helping petals to invade nutrient-rich regions of the Petri dish. By contrast, ACD or excessive RCD can inhibit growth, suggesting a balance between the altruistic and destructive effects of RCD.

## 2 Mathematical model and experimental method

### 2.1 Mathematical model

We extend the reaction–diffusion model (1.1) to include both accidental and regulated cell death. Adopting the notation used in (1.1), we follow Tam et al. [14] and investigate the nutrient-limited growth of a yeast colony biofilm on a Petri dish, and neglect the extracellular matrix as well as any mechanical effects. We assume that the mechanisms of ACD and RCD each depend exclusively on the local dimensional nutrient concentration, *g*^*^. Deviating from Tam et al. [14], we define the dimensional density of dead cells having undergone ACD to be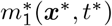. In the model, ACD occurs when the nutrient concentration is sufficiently low. Specifically, ACD occurs when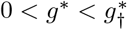, where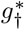is a constant nutrient threshold below which ACD occurs. We assume also that the number of cells dying by ACD in unit time is proportional to the product of the dimensional living-cell density, *n*^*^ and the difference in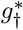 and *g*^*^, having dimensional ACD rate constant,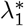.

We assume that regulated cell death (RCD) occurs when the nutrient concentration *g*^*^ satisfies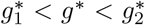. The constants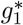and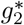 define the window within which RCD can occur. Similar to ACD, we define the dimensional RCD cell density to be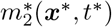 and assume also that the number of cells dying by RCD is proportional to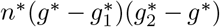, where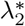 is the dimensional RCD rate constant. As time evolves, we assume that ACD and RCD cells breakdown with respective rates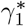and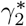. A proportion of these broken down cells release nutrients for further consumption. We define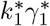and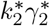to be the amount of nutrient released from dead cells, per unit area, in unit time.

Given these assumptions, the coupled four-species reaction–diffusion model is:

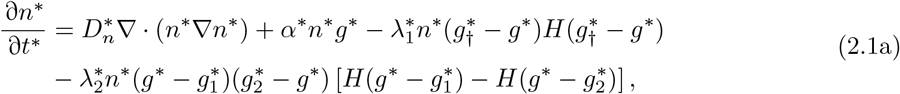

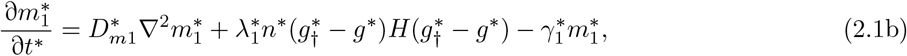

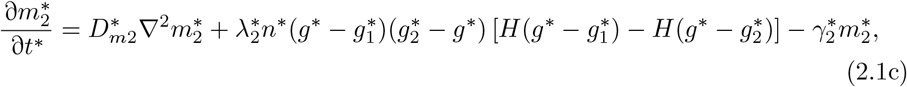

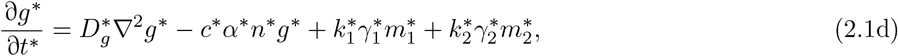

where *H*(·) is the Heaviside step function and ∇^2^ is the spatially two-dimensional Laplace operator. The constants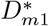and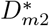are the respective dimensional diffusivities of ACD and RCD cells.

To close the model (2.1), we require appropriate initial and boundary conditions. Since the dynamics occur within a solid Petri dish, we impose the following no-flux conditions at the boundary *∂S*^*^:

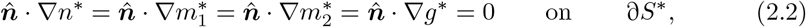

Where 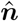 is the outward pointing unit normal to ∂*S*^*^. We assume that initially there are no dead cells present, and that the nutrient concentration is spatially uniform. The initial conditions are then:

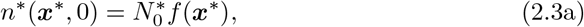

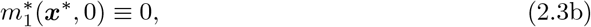

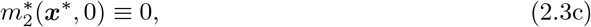

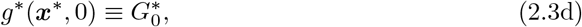

where *f* is a dimensionless function controlling the initial profile of the living cells, and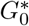 is the initial nutrient concentration. The governing equations (2.1) and conditions (2.2) and (2.3) define a system of four coupled partial differential equations in terms of four unknown quantities: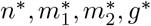.

#### 2.1.1 Nondimensionalisation

We nondimensionalise the system (2.1)–(2.3) by introducing the scalings:

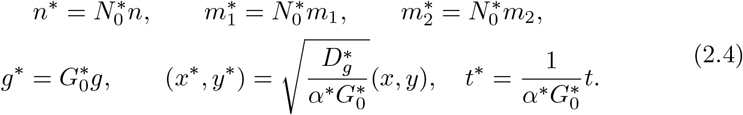

Here cell densities and the nutrient concentration have been scaled by the initial cell density and nutrient concentration,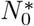and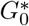, respectively. Spatial co-ordinates have been scaled with respect to the ratio between nutrient diffusion and the rate of nutrient uptake, and time has been scaled with respect to the reciprocal of the nutrient uptake rate. Substituting the scalings (2.4) into (2.1), we obtain the dimensionless system:

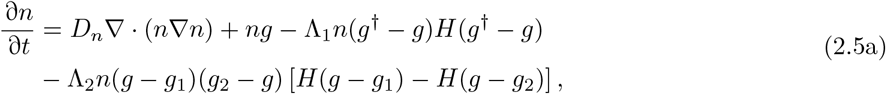

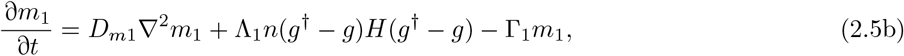

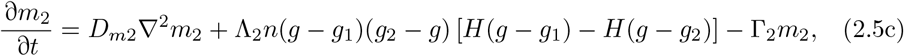

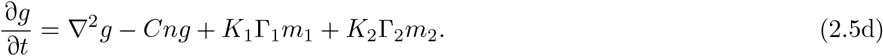

The dimensionless system (2.5) involves the thirteen dimensionless groups:

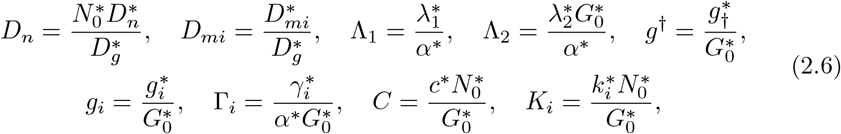

where *i* ∈ (1, 2) and ∇^2^ is now taken as the dimensionless Laplace operator. The constants *D*_*n*_ and *D*_*mi*_ are the ratios of the living, ACD and RCD cell diffusivity to nutrient diffusivity respectively. The groups Λ_1_ and Λ_2_ are respective dimensionless ACD and RCD rates. The parameters *g*^†^ and *g*_*i*_ are dimensionless nutrient thresholds. The breakdown/decay rates Γ_*i*_ represent the respective ratios of ACD and RCD breakdown rates to the nutrient uptake rate. The constant *C* is the dimensionless consumption rate, and the *K*_*i*_ are dimensionless measures of the quantity of nutrient released when an ACD or RCD cell breaks down. From (2.2) and (2.3), the dimensionless boundary and initial conditions to be applied to (2.5) are:

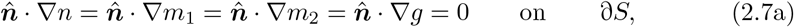

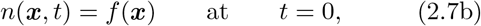

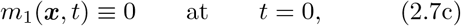

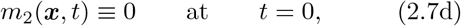

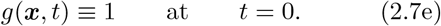

Equations (2.5) and (2.7) then form the closed dimensionless model that we investigate numerically.

#### 2.1.2 Numerical Methods

We obtain numerical solutions to the dimensionless system (2.5) and (2.7) in one and two spatial dimensions using the method of lines. Spatial derivatives are approximated using second-order accurate differentiation matrices, while temporal derivatives are discretised using the first-order Euler method. In all spatially one-dimensional simulations, we chose the initial profile of the dimensionless living-cell density to be *f* (*x*) = exp(− *x*^2^). Since this choice for *f* is even, and the system (2.5) contains only even derivatives in *x, y*, and the no-flux conditions (2.7a) are all homogeneous, it follows that solutions for *n, m*_1_, *m*_2_ and *g* are symmetric about *x* = 0 in this spatially one-dimensional case. In all spatially two-dimensional solutions, we used initial conditions that also exhibit this symmetry. Hence, in both the spatially one and two-dimensional cases, it is convenient to solve the problem numerically on *x >* 0, imposing symmetry conditions whereby the first derivative of each of the dependent variables vanishes at *x* = 0 and *y* = 0.

### 2.2 Experimental Method

In this section we describe the experimental method used to obtain the results presented in Figure 1.1. In our experiments, three different varieties of yeast were used: Σ1278b, a diploid prototrophic strain of *Saccharomyces cerevisiae*; W303a (*MAT* a *leu2-3*,*112 trp1-1 can1-100 ura3-1 ade2-1 his3-11*,*15 ssd1-d*), and an in-house isolate of *Magnusiomyces magnusii*. Yeast Peptone Dextrose (YPD) medium (10 g L^−1^ yeast extract, 20 g L^−1^ peptone, 20 g L^−1^ glucose) with 0.6% agar was prepared by filter sterilising 2 x YPD

and mixing with an equal volume of molten 12 g L^−1^ agar. Phloxine B was added at 10 µm. 5 mL yeast cultures were grown for 48 hours in liquid YPD prior to inoculation to plates which was either as a 5 µL spot in the centre of medium in a 90 mm round Petri dish or as a streak, applied with a 1 µL plastic inoculation loop to medium in a 100 mm square Petri dish. Plates were cling wrapped and incubated agar side down for 10 days at 25 °C. Macroscopic plate images were captured with an Apple iPhone 12 Pro and microscopic imaging. For experiments with W303a cells were grown overnight at 30°C in YPD and colonies were grown from single cells for a period of 5 days on YPD agar plates containing 10 µm Phloxine B before images were taken using a Leica MZFLIII dissecting microscope at 10 × magnification under GFP illumination (Ex488/Em520) and captured with a Cairn Scientific CellCam 200CR using ImageJ software.

## 3 Numerical results and discussion

We solve the system of equations (2.5) and (2.7) numerically in one and two spatial dimensions to investigate how ACD, RCD, cell breakdown and nutrient release influence colony biofilm growth dynamics. We show that our mathematical model is capable of reproducing qualitatively the three experimental patterns of death and morphology observed in Figure 1.1. The model builds on our experimental results, hypothesising that death in the colony centre is a consequence of ACD, and death near the proliferating front is a consequence of RCD.

### 3.1 Spatially one-dimensional numerical results

Here we present spatially one-dimensional numerical solutions of (2.5) and (2.7). Our results suggest that the biofilm will grow into a state in which ACD cells are found primarily in the core of the colony (*i*.*e*. a necrotic core) and that RCD cells can appear in an annulus following the colony front, which then forms a plausible explanation of the patterning observed in Figure 1.1.

#### 3.1.1 Colonies with ACD and no nutrient release can form a necrotic core

We begin by considering the case by which ACD is the only present cell-death mechanism, and examine the effect that increasing the rate of ACD has on the expansion of the colony front in the absence of breakdown (Γ_1_ = 0). In the absence of cell death (Λ_1_ = 0), our numerical method reproduces numerically (see Figure 3.1) the travelling-wave solutions obtained by Tam et al. [14]. Introducing ACD by setting Λ_1_ *>* 0 changes the colony composition significantly. Rather than occupying the entire colony, living cells now only appear in a pulse close to the leading edge. Behind the front, the living-cell density decays, and the density of the ACD cells increases, as Figure 3.1 shows. This scenario represents a front of proliferating cells surrounding a large necrotic core, as observed in Figure 1.1A. As expected, increasing the ACD rate Λ_1_ decreases the expansion speed of the colony. This effect occurs because nutrient consumption and cell proliferation drive yeast-colony expansion. Reinforcing this, the amplitude of the living-cell pulse monotonically decreases as the ACD rate Λ_1_ is increased. Without subsequent ACD breakdown and nutrient release, the ACD-cell density settles to the constant value 1*/C*. For small Λ_1_, living cells are capable of consuming all of the nutrient at the centre of the colony. If the rate of ACD is sufficiently large, for example Λ_1_ = 1, we find that the cells die before all of the nutrient in consumed.

**Fig. 3.1:**
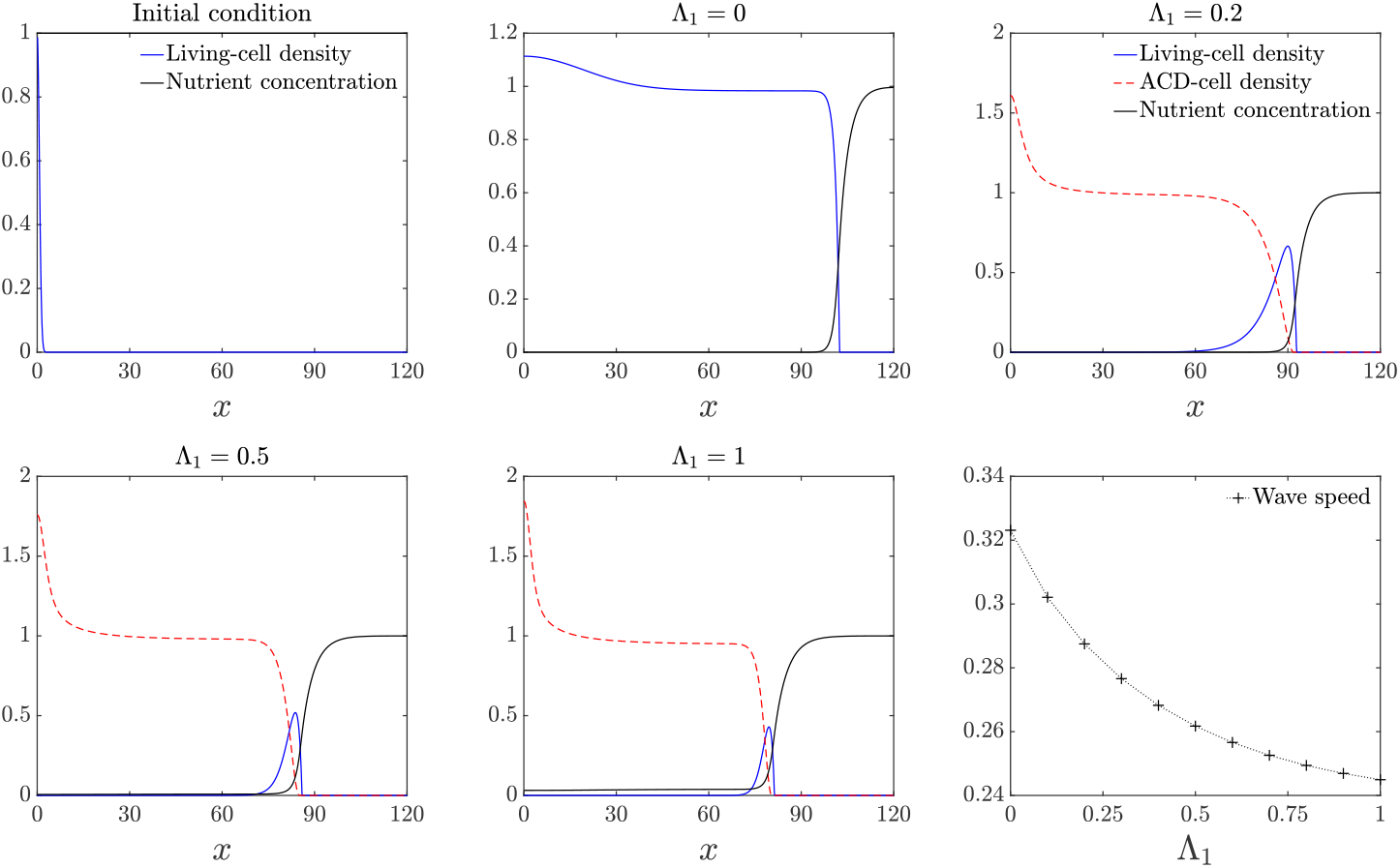
Numerical solutions for the living-cell density *n*, ACD-cell density, *m*_1_, and nutrient concentration *g*, plotted against *x* for increasing Λ_1_. The top-left panel indicates the initial condition. The bottom-right panel shows the expansion speed plotted against Λ_1_. The remaining panels indicate solutions at *t* = 300. The parameter values used used to obtain these numerical results are given by: *D*_*n*_ = 0.47, *D*_*m*1_ = 0.001, *C* = 1, Γ_1_ = 0, *g*_†_ = 0.25.

We now consider the effect of cell breakdown and nutrient release for the case in which ACD is the only present cell-death mechanism. As we see in Figure 3.2, for sufficiently small values of the ACD nutrient-release parameter *K*_1_, the pulse profile for the living cells that occurs with no nutrient release is preserved. As *K*_1_ increases, the pulse increases in amplitude and width. Expansion speed also increases approximately linearly with the amount of nutrients released *K*_1_. Increasing *K*_1_ provides the colony with more nutrient, facilitating more cell proliferation, increased cell density, and faster expansion. Unlike in Figure 3.1, the ACD-cell density takes a pulse profile, due to the breakdown of cells behind the front. For sufficiently large *K*_1_, enough nutrient is released to the colony that the living-cell density transitions to a wave-front profile, such that living cells, ACD cells, and nutrient are all present throughout the colony.

#### 3.1.2 Colonies with ACD, RCD, and nutrient release can have death in an edge-trailing ring and in the core

We now investigate the phenotype in Figure 1.1B, where there is a core of dead cells, and a distinct ring of dead cells trailing the front. We show that introducing RCD at a higher rate than ACD can give rise to this ring structure. We consider the case in which the rate of ACD breakdown and nutrient release is much smaller than the corresponding RCD rate. As the rate of RCD breakdown increases, the rate at which nutrient is released back to the colony is increased, speeding its expansion. Examining Figure 3.3, in contrast to Figures 3.1 and 3.2, a dominant RCD rate means that the living cells occupy the entire domain. Since ACD is low, living cells can persist in the core where nutrient concentration is low. Although the number of ACD cells is small overall, they are maximised in the colony centre (*x* = 0), and decrease approximately linearly away from the centre. On the other hand, RCD occurs with maximum rate at an intermediate nutrient concentration. When RCD cells break down, the distribution of RCD cells takes the form of a pulse following the leading edge, whose amplitude is modulated by the RCD breakdown rate, Γ_2_. Under the assumption of radial symmetry, this would appear as a red ring following closely behind the front, and so could be representative of the experimental result presented in Figure 1.1B. In this experiment, there is a less-pronounced region of dead cells in the centre of the colony, which these solutions predict to be ACD cells. If the RCD breakdown rate Γ_2_ is small, the ratio of RCD to ACD cells increases, because small Γ_2_ increases the amplitude of the RCD pulse. As the colony expands, the nutrient concentration decays, meaning that the rate of RCD is reduced. Since the rate of RCD is significantly larger than ACD, the colony is able to maintain life in the core. Interestingly, the bottom-right panel of Figure 3.3 demonstrates that even small RCD breakdown rates yield comparatively large increases in expansion speed. We hypothesise that the ring phenotype in Figure 1.1B might occur due to fast RCD compared to ACD and a sufficiently low rate of nutrient release. Even a small amount of nutrient release would be sufficient to increase colony expansion speed, and would correspond to the most pronounced cell-death ring pattern trailing the leading edge.

**Fig. 3.2:**
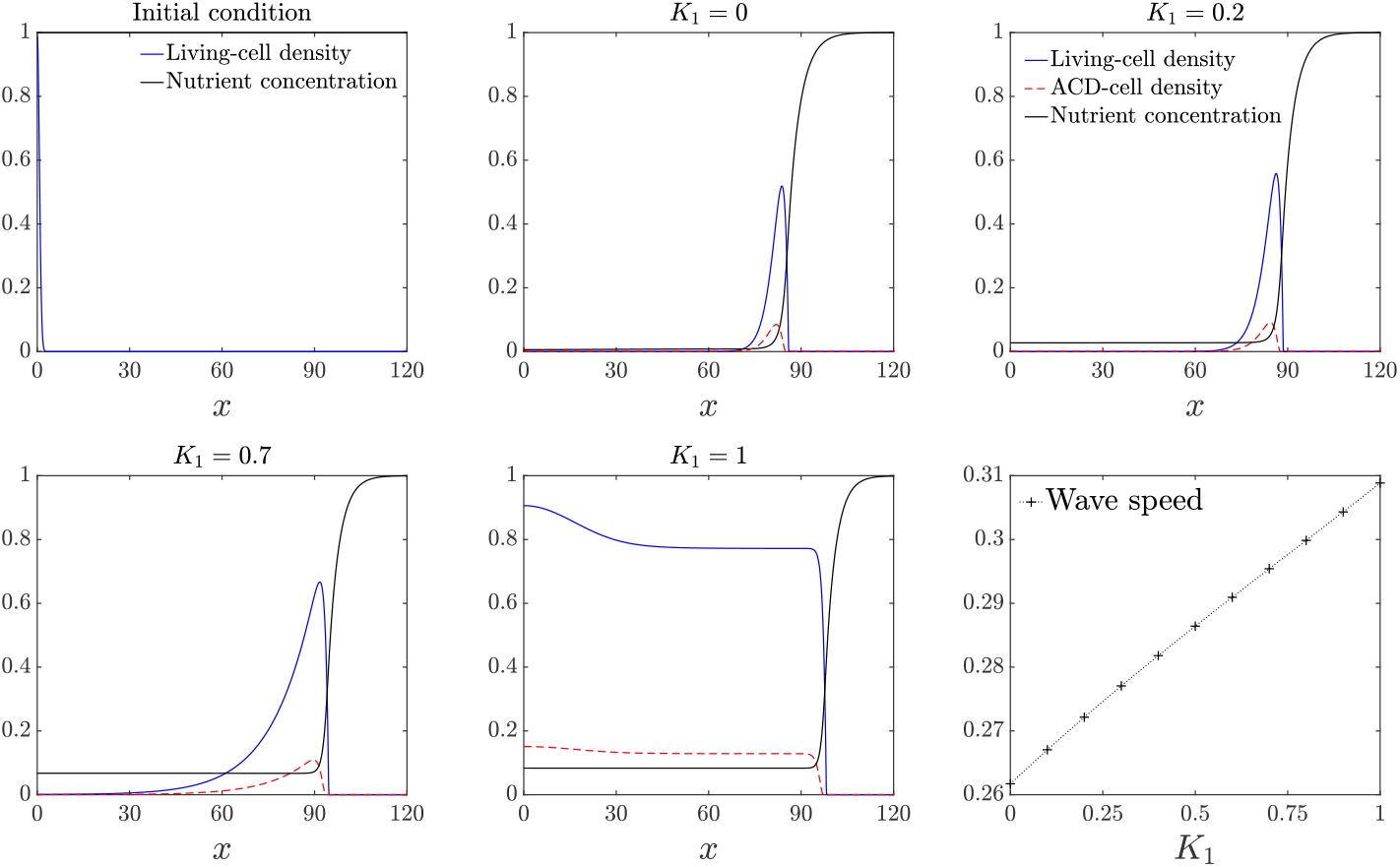
Numerical solutions for the living-cell density *n*, ACD-cell density, *m*_1_, and nutrient concentration *g*, plotted as a function of *x* for increasing *K*_1_. The top-left panel indicates the initial condition. The bottom-right panel shows the expansion speed plotted against *K*_1_. The remaining panels indicate solutions at *t* = 300. The parameter values used for this simulation are: Λ_1_ = 0.5, *D*_*n*_ = 0.47, *D*_*m*1_ = 0.001, *C* = 1, Γ_1_ = 0.5, *g*_†_ = 0.25.

**Fig. 3.3:**
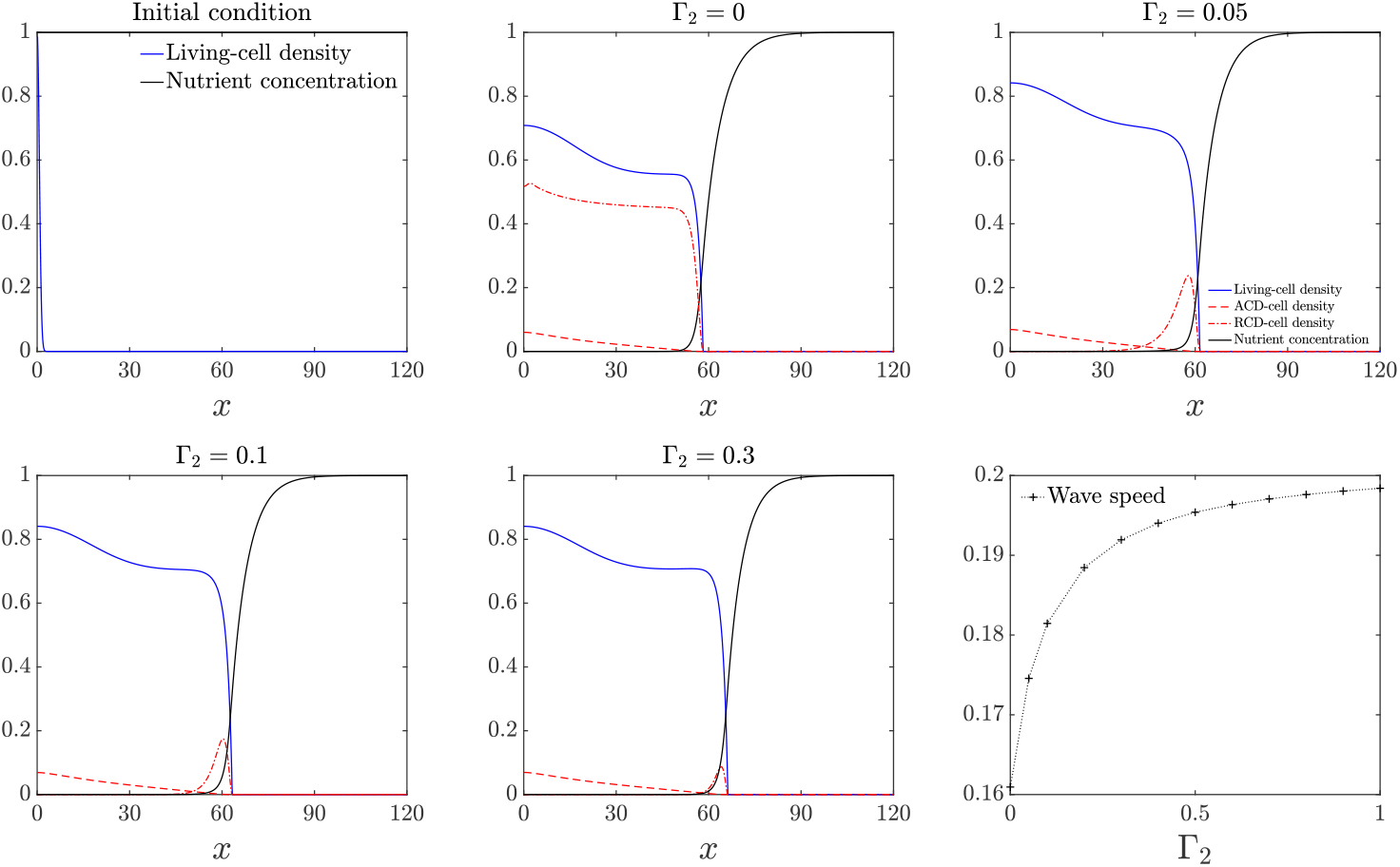
Numerical solutions for the living-cell density *n*, ACD-cell density, *m*_1_, RCD-cell density, *m*_2_, and nutrient concentration, *g*, plotted at *t* = 300 against *x* with *K*_1_ = 0, *K*_2_ = 0.5 and increasing Γ_2_. The top-left panel indicates the initial condition. The bottom-right panel shows the expansion speed plotted against Γ_2_. The remaining parameter values used for this simulation are: *D*_*n*_ = 0.47, *D*_*m*1_ = 0.001, Λ_1_ = 0.001, Λ_2_ = 0.5, *C* = 1, Γ_1_ = 0, *g*_†_ = 0.25, *g*_1_ = 0, *g*_2_ = 1.

### 3.2 Spatially two-dimensional numerical results

The spatially one-dimensional solutions in Section 3.1 apply to the experiments of Figure 1.1A–B, where the patterns exhibit approximate radial symmetry. In the absence of cell death (*m*_1_ ≡ *m*_2_ ≡ 0), it has previously been shown that planar travelling-wave solutions to the model (2.5) can be linearly unstable to transverse sinusoidal perturbations [14, 36]. We now investigate numerically the spatiotemporal stability of the spatially one-dimensional solutions obtained in Section 3.1 to qualitatively understand how cell death affects colony morphology. We use the spatially one-dimensional numerical solutions to (2.5) and (2.7) at *t* = 300 as the base states for our numerical linear stability analysis, and denote these solutions respectively as: *n*_0_, *m*_10_, *m*_20_, and *g*_0_. We then use transversely perturbed versions of these solutions as initial conditions to spatially two-dimensional numerical solutions of (2.5) and (2.7), and observe how the perturbations evolve over time. Formally, the initial conditions used for the two-dimensional simulations are:

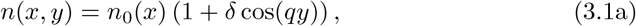

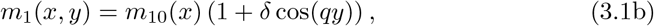

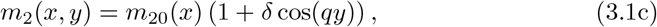

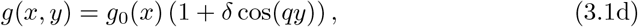

where 0 *< δ*≪ 1 is the initial amplitude of the perturbation, and *q >* 0 is the wavenumber. Planar front solutions to (2.5) without cell death can be unstable to long-wavelength (small *q*) transverse perturbations, depending on the ratio of cell diffusivity to nutrient diffusivity [36]. In all solutions, we use *δ* = 0.1 and *q* = 6*π/*50 = 0.3770, which is close to the most unstable wave number for *D* = 0.47, determined by Tam et al. [14]. In Figure 3.4A, we reproduce numerically the unstable floral morphology with these parameters in the absence of cell death.

**Fig. 3.4:**
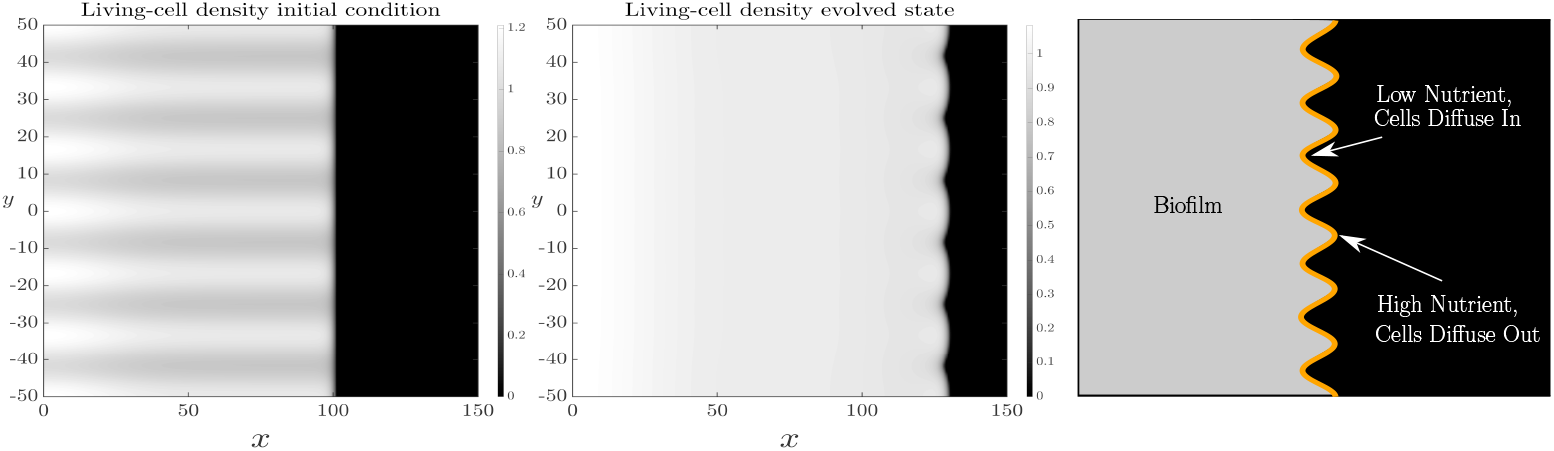
Spatially two-dimensional numerical solution of (2.5) and (2.7a) subject to the transversely perturbed initial condition (3.1) on the domain *x* ∈ (0, 150), *y* ∈ (−50, 50) in the absence of cell death. The first panel shows the initial condition, the centre panel shows the corresponding solution at *t* = 100, and the final panel is a schematic illustrating the instability mechanism. The specific parameter values that have been used are: *D*_*n*_ = 0.47, *C* = 1, Λ_1_ = 0, Λ_2_ = 0, *δ* = 0.1, *q* = 6*π/*50.

Figure 3.4B illustrates the mechanism of the instability. As Müller and van Saarloos [36] explain, when the front is perturbed there is a trade-off between nutrient supply, which amplifies the perturbation, and cell diffusivity, which smooths the perturbation. When the colony protrudes ahead of the undisturbed front, it enters a comparatively nutrient-rich region, which aids cell proliferation and further protrusion. Nutrients are more depleted behind the undisturbed front, inhibiting cell proliferation there. Cell diffusion counteracts the amplifying effect of nutrient supply [36]. The cellular diffusive flux is proportional to −*n*∇ *n*. When the colony protrudes ahead of the undisturbed front, cell diffusion will oppose the protrusion and smooth the perturbation. Conversely, if there are unoccupied regions behind the undisturbed front, cells will diffuse towards these regions. Due to these competing effects, increasing the diffusivity ratio *D*_*n*_ suppresses the instability for shorter wavelength modes [36], for which the lengthscale required for cell diffusion to stabilise the front is shorter. For sufficiently large *D*_*n*_, the instability is completely eliminated for all modes [36]. Using spatially two-dimensional numerical solutions, we will investigate qualitatively how cell death influences these instability dynamics. Since the growth rates of these transverse perturbations are small [36], we explore the instability numerically, solving until *t* = 100 beyond the linear regime.

#### 3.2.1 Cell death amplifies floral pattern formation

Cell death affects the balance between cell diffusion and nutrient diffusion, and hence the instability mechanism described in Section 3.2. Spatially two-dimensional numerical solutions with cell death present support this idea. Figure 3.5 shows spatially two-dimensional solutions of (2.5), (3.1) and (2.7a), for the case in which ACD is the only present cell-death mechanism, and in the absence of cellular breakdown and nutrient release. The results indicate the stability of the spatially one-dimensional numerical results presented in Figure 3.1. Qualitatively, it is observed that ACD in isolation amplifies the instability, having increased petal protrusion compared to Figure 3.4, where there was zero cell death. With ACD is present, but does not lead to nutrient release, the amplitude of the living-cell density declines with increasing Λ_1_, as Figure 3.1 shows. Smaller cell density suppresses the stabilising impact of cell diffusion. Since there is no nutrient release, the destabilising effect of the nutrient concentration is unchanged. The net result is an increase in petal amplitude.

**Fig. 3.5:**
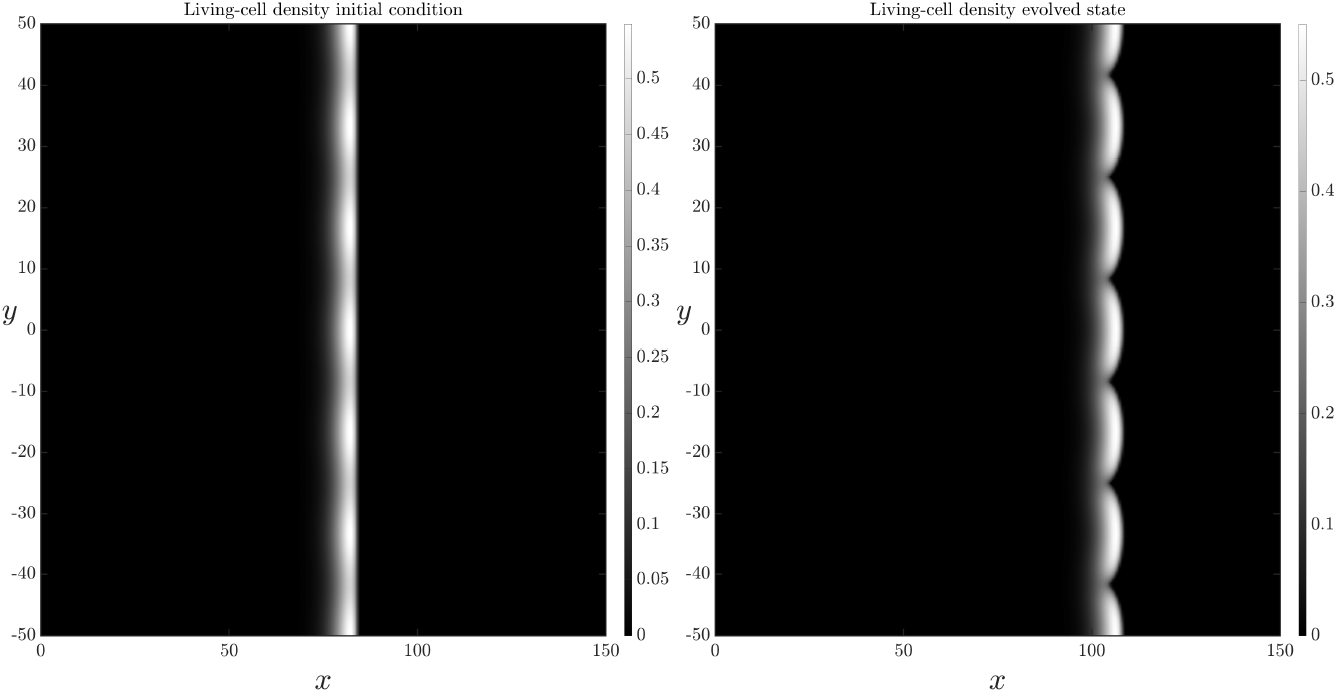
Spatially two-dimensional numerical solution of (2.5) and (2.7a) subject to the transversely perturbed initial condition (3.1) on the domain *x* ∈ (0, 150), *y* ∈ (−50, 50). There is ACD, but no nutrient release or RCD. The left-hand panel shows the initial condition. The right-hand panel shows the solution at *t* = 100. The parameter values that have been used are: *D*_*n*_ = 0.47, *C* = 1, Λ_1_ = 0.5, Λ_2_ = 0, Γ_1_ = 0, *K*_1_ = 0.

With nutrient release from ACD-cell breakdown, nutrient is resupplied to the colony, enabling more cell proliferation, and hence increasing the living-cell density. This was discussed in Section 3.1 and Figure 3.2. The cell density remaining higher neutralises the suppression of cell diffusion discussed above. Therefore, nutrient release tends to lessen the floral instability. This effect occurs in Figure 3.6, where petal amplitude is similar to, but smaller than, the petal size with ACD only and no nutrient release (Figure 3.5).

**Fig. 3.6:**
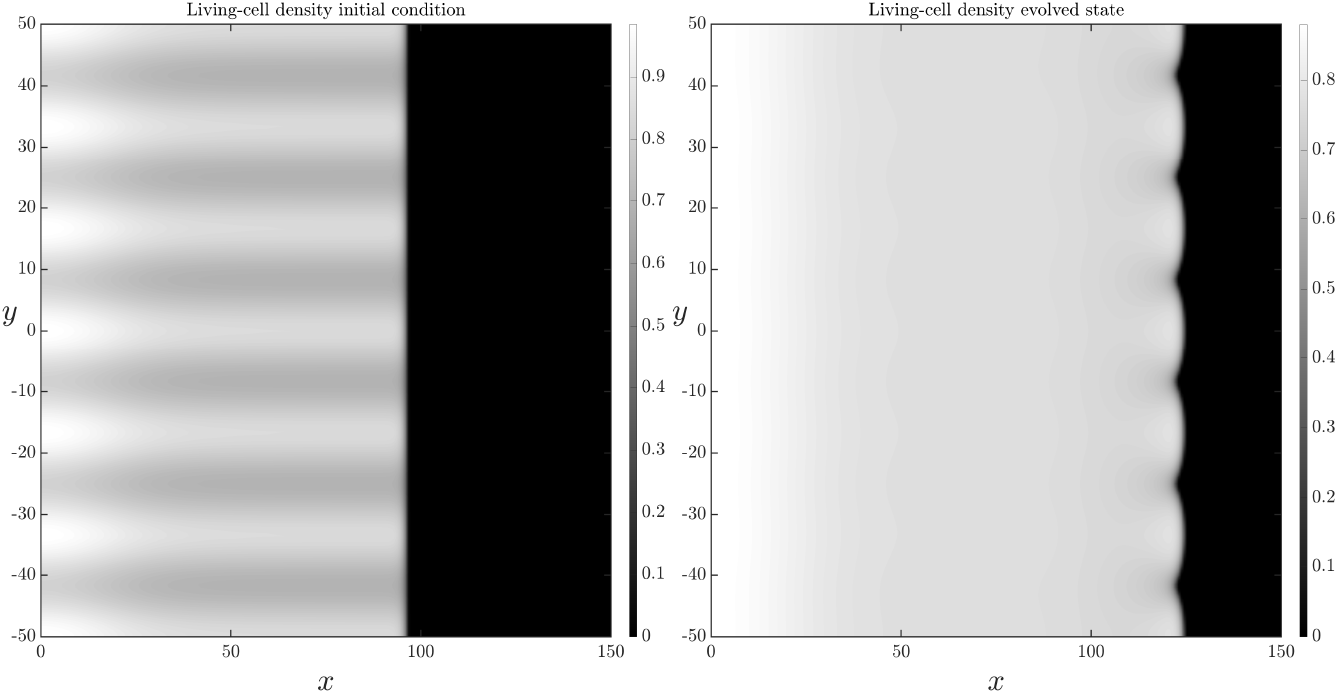
Spatially two-dimensional numerical solution of (2.5) and (2.7a) subject to the transversely perturbed initial condition (3.1) on the domain *x*∈ (0, 150), *y*∈ (−50, 50). There is ACD and nutrient release, but no RCD. The left-hand panel shows the initial condition. The right-hand panel shows the solution at time *t* = 100. The parameter values that have been used are: *D*_*n*_ = 0.47, *C* = 1, Λ_1_ = 0.5, Λ_2_ = 0, Γ_1_ = 0.5, *K*_1_ = 1.

In Section 3.1 it was shown that a combination of ACD and RCD capable of breakdown and nutrient release can produce the spatial patterns observed in Figure 1.1B. We now explore numerically how these features influence the instability by perturbing the solutions presented in Figure 3.3. With the low rate of nutrient release that yielded the ring, Figure 3.7 shows that the interface remains unstable. However, the combination of ACD and RCD decreases the petal amplitude. This reduction in amplitude is due to the fact that RCD and nutrient recovery occurs near the proliferating rim, promoting cell proliferation there. Cell proliferation increases the living-cell density, inhibiting the floral instability. Our results suggest that the primary function of RCD and nutrient release is to increase the overall colony expansion speed, and not necessarily enhance petal formation. In less favourable conditions, the floral morphology might help the yeast colony to more easily invade previously unoccupied regions, and increase overall nutrient supply to the cells. Nutrient release from dead cells improves the growth conditions near the leading edge. Therefore, there is less need for amplified petal formation as the colony expands, even though the instability persists.

**Fig. 3.7:**
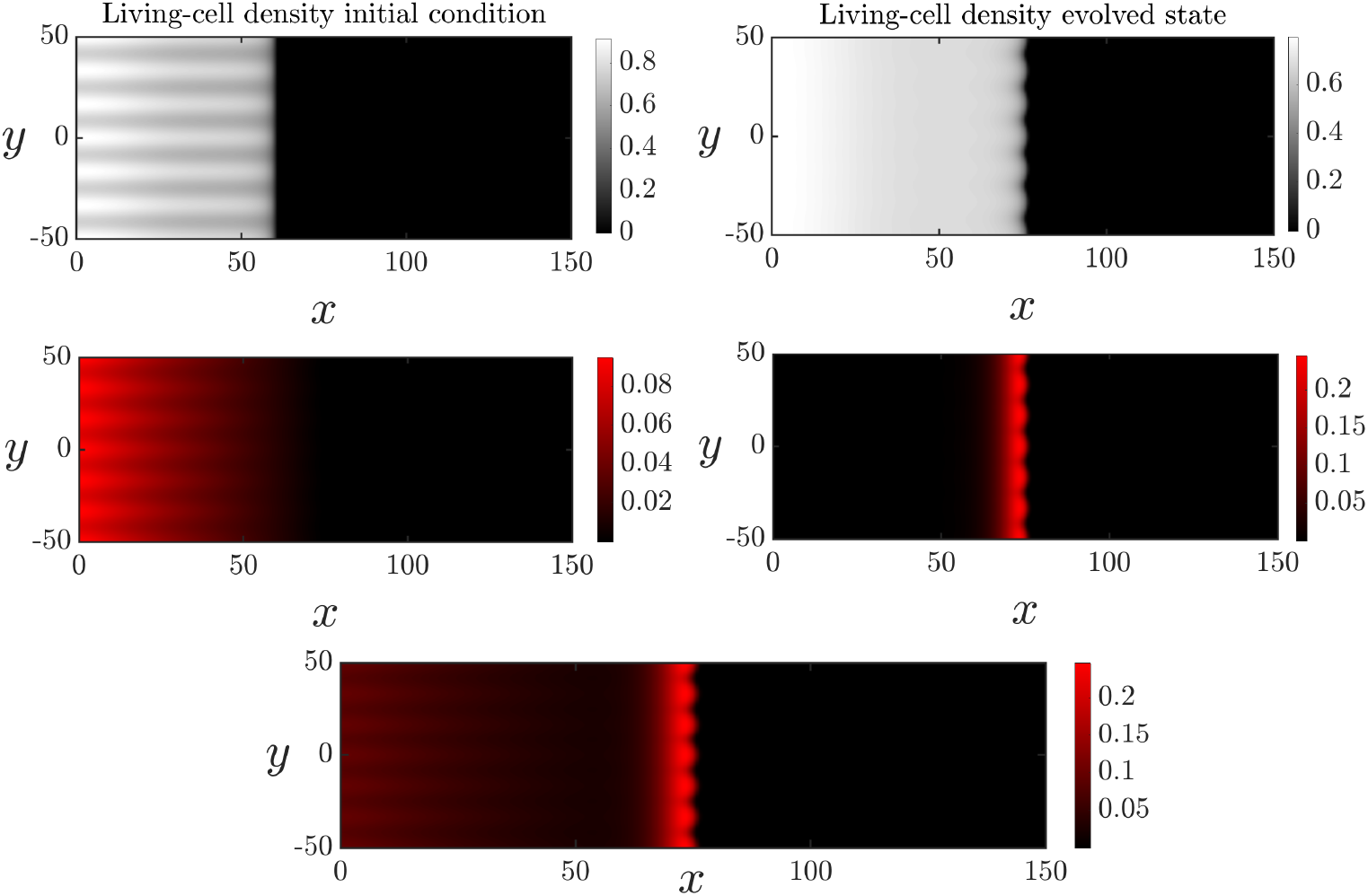
Spatially two-dimensional numerical solution of (2.5) and (2.7a) subject to the transversely perturbed initial condition (3.1) on the domain *x* ∈ (0, 150), *y* ∈ (− 50, 50). There is ACD, RCD and nutrient release. The top row shows the living-cell density in its initial (left) and evolved (state). The middle panel shows the ACD-cell density (left) and RCD-cell density (right) in their evolved states. The bottom panel shows the sum *m*_1_ + *m*_2_ of the dead cell densities. The parameter values that were used are: *D*_*n*_ = 0.47, *C* = 1, Λ_1_ = 0.001, Λ_2_ = 0.5, Γ_1_ = 0, Γ_2_ = 0.05, *K*_2_ = 0.5.

#### 3.2.2 Solutions with linear cell diffusion do not develop floral morphology

Previous works have shown that nonlinear cell diffusion is required in order for planar-front solutions of our model (2.5) in the absence of cell death to be unstable [36], and that planar fronts are stable with linear cell diffusion [58]. Here we demonstrate that this is also the case when cell death is present. To do this, we solved the system (2.5) with linear, Fickian cell diffusion. This modified system was then solved with the effects of ACD, RCD, and nutrient release present. The corresponding numerical results are shown in Figure 3.8, where it is observed that linear diffusion results in an apparently stable planar front. Therefore, although cell death can amplify the petal-forming instability, nonlinear cell diffusivity is the instability mechanism. Since yeast cells are immotile, living-cell diffusion is a phenomenological description for how cells colonise the Petri dish. Nonlinear cell diffusivity has been justified because it yields solutions with compactly-supported cell density, as occurs in experiments. Nonlinear diffusivity might also arise due to random movement of cells with aspect ratios not equal to unity [59], and also because the diffusive flux (spread of cells) increases with elevated cell density, in addition to the cell-density gradient. However, since cell diffusivity is phenomenological, the most suitable cellular diffusivity that corresponds to experiments is unknown. In Figure 1.1, the colonies in Panels A and B have little floral pattern formation, whereas the colony in Panel C does have a floral pattern. Since nonlinear cell diffusivity predicts slow-growing instability, we cannot determine whether the experiments in Panels A and B are too early to observe the instability, or whether the instability will never develop. Both Figures 3.7 and 3.8 indicate the ring of cell death. Therefore, the broad impact of ACD and RCD on the distribution of living and dead cells is independent of the choice of cellular diffusivity.

**Fig. 3.8:**
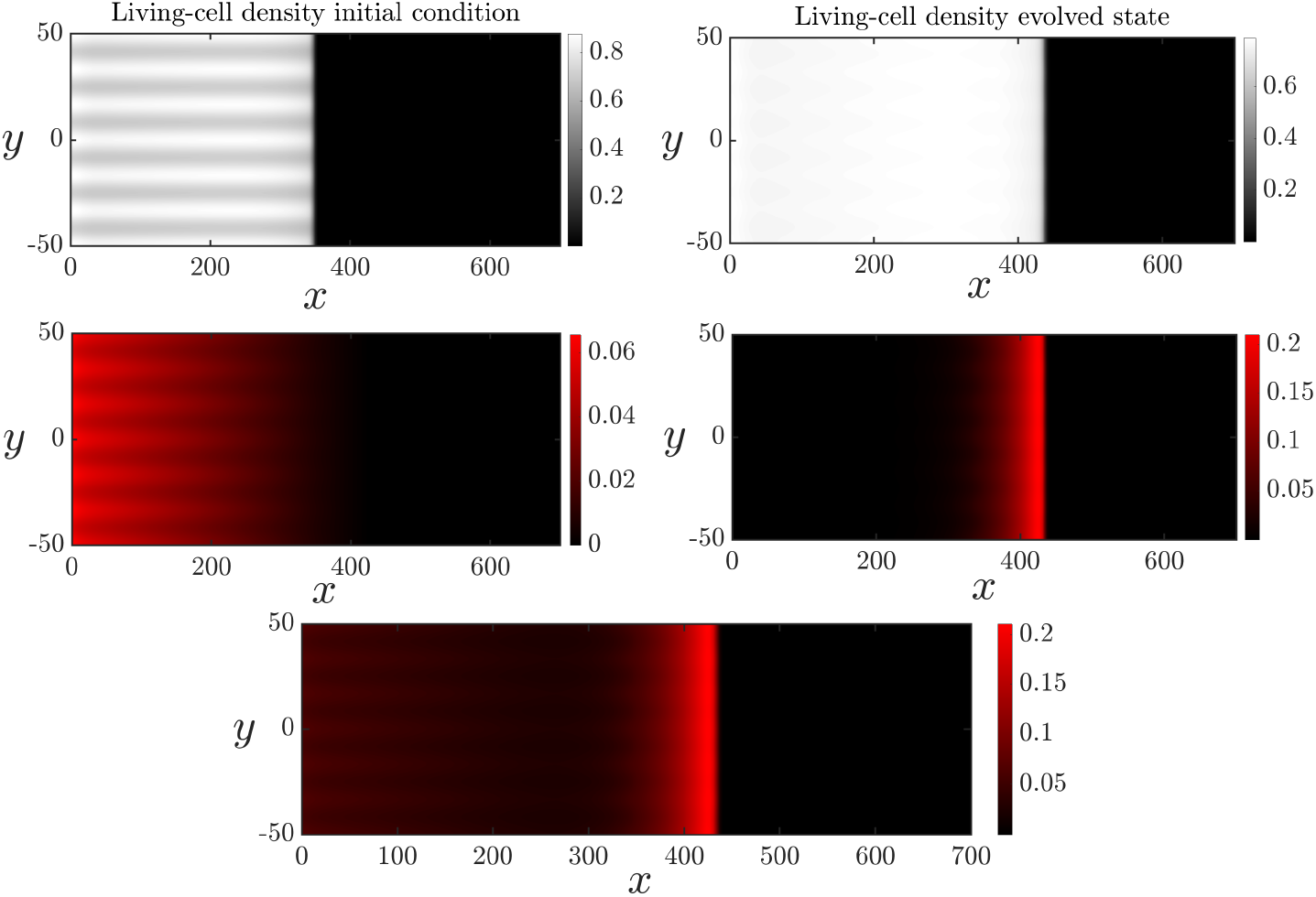
Spatially two-dimensional numerical solution of (2.5) and (2.7a) subject to the transversely perturbed initial condition (3.1) on the domain *x* ∈ (0, 700), *y* ∈ (−50, 50) for the case of linear diffusion. The parameters were chosen to match Figure 3.7

## 4 Conclusions

Yeasts can alter their growth to adapt to different environments. These adaptations help yeasts to survive in both natural environments and controlled laboratory experiments. In this article, we have used experiments and mathematical modelling to investigate the effects of accidental cell death (ACD) and regulated cell death (RCD) on the nutrient-limited growth of yeast colony biofilms.

We used the vitality dye Phloxine B to indicate regions of elevated cell death in yeast colony biofilms. Our experiments reveal three patterns of cell death. One pattern involves a colony that develops a region of cell death at the centre, with living cells being found closer to the proliferating rim. In a different experiment, the most pronounced region of cell death occured in a ring that trails the leading edge. This experiment was also replicated in a rectangular Petri dish, where we observed a region of cell death trailing the front, which evolves into a non-uniform spatial pattern. Our objective was to develop a mathematical model that could explain the three patterns of cell death observed in experiments.

Distinguishing between ACD and RCD helps to formulate a potential explanation for the different colony patterns observed in Figure 1.1. ACD occurs in harsh environments, where the cell is unable to sustain life. RCD occurs when a cell would be able to survive, but instead elects to die in response to an environmental cue. RCD is altruistic, where the death benefits the colony as a whole. The benefit to the colony is hypothesised here as being due to the release of nutrients back to the colony. Nearby cells can then consume the returned nutrients, spurring increased cell proliferation and colony expansion.

In this work, we have extended an existing reaction–diffusion model for the expansion of yeast colony biofilms [14, 36] to incorporate the effects of ACD and RCD. This extended system involves four coupled nonlinear partial differential equations, which collectively describe the spatiotemporal evolution of the living cells and nutrient concentration within the colony, as well as two species of dead cells: those that have died by either ACD or RCD. Numerical solutions of this extended system were shown to reproduce qualitatively the patterning observed in our experiments. In the absence of RCD, we used our model to demonstrate that the colony can form a necrotic-core-like composition, with living cells mostly being found close to the leading edge. If RCD is introduced, the colony takes a more complex composition. In this case, living cells will, in general, persist throughout the colony. Cells that die by ACD are most abundant in the centre, and cells dying by RCD occur in a pulse close to the leading edge. This behaviour is reminiscent of the experimental ring phenotype in Figure 1.1B. Varying model parameters demonstrated that the breakdown of RCD cells together with nutrient release can increase the speed of expansion. The impact on expansion speed is greatest at low amounts of RCD breakdown, whereas large amounts of breakdown incur diminishing returns. Our model suggests that even small amounts of RCD can be impactful for colony expansion.

Numerical solutions in two spatial dimensions enabled us to investigate the non-uniform front patterns observed in our rectangular experiments Figure 1.1C. Previous work by Müller and van Saarloos [36] shows that a reaction–diffusion model without cell death could predict petal-like structure formation in yeast colony biofilms [14]. These petals are advantageous for colony growth, enabling regions of the colony boundary to invade unoccupied regions faster. Our results demonstrate that, although cell death slows overall expansion, it enhances petal formation. This enhanced petal formation could be a secondary advantage to the colony. Like previous mathematical models [36, 56], nonlinear cell diffusion was necessary for our model to give rise to spatially unstable numerical solutions.

Careful interpretation of our results requires considering some limitations to this study. Introducing ACD and RCD to existing models for nutrient-limited colony biofilm growth generates a system of four coupled reaction–diffusion equations. It also requires introducing new parameters, specifically, the rate of ACD and RCD, the rate of cellular breakdown and nutrient release, the proportion of nutrient that can be re-consumed, and the nutrient concentrations thresholds at which ACD and RCD occur. The results presented in this paper are only a subset of the possible behaviours, and depend on the choices of parameter values. Furthermore, previous minimal reaction–diffusion systems were amenable to travelling-wave analysis and linear stability analysis. Since each reaction–diffusion equation is second-order, performing a travelling-wave analysis of the model with cell death would involve the complicated task of analysing an eight-dimensional dynamical system. Without this analysis, we do not have theoretical predictions of the expansion speed or the growth rate of perturbations in the two-dimensional solutions, and instead rely on numerical solutions to analyse the model qualitatively. Another limitation is that the Phloxine B dye cannot distinguish between accidental and regulated cell death. Although experimentalists believe that the trailing ring of cell death is due to regulated cell death, we are unable to confirm this definitively. Consequently, the key contribution of this work is using the mathematical model to generate a theoretical hypothesis for how RCD might be responsible for the observed cell-death patterns. Future experimental work could focus on isolating the nature of cell death in either the core or the outer ring.

## Acknowledgements

We acknowledge funding from the Australian Research Council (Grant numbers DP230100406, and DE240100897).

## Authorship Contribution Statement

- **DJN:** Methodology, software, validation, formal analysis, investigation, writing — original draft, writing — review & editing, visualisation.
- **AKYT:** Methodology, writing — original draft, writing — review & editing, supervision, funding acquisition.
- **JEFG:** Conceptualization, methodology, writing — review & editing, supervision, funding acquisition.
- **TK:** Investigation, data curation.
- **JMG:** Investigation, data curation, writing — original draft, supervision.
- **CWG:** Conceptualization, methodology, investigation, data curation, writing — original draft, writing — review & editing, writing — review & editing, funding acquisition.
- **VJ:** Conceptualization, writing — review & editing, supervision, funding acquisition.
- **BJB:** Conceptualization, methodology, writing — review & editing, supervision, project administration, funding acquisition.

## Data Availability

Numerical code is available on request.

